# Multi-cancer classification; an analysis of neural network complexity

**DOI:** 10.1101/2022.01.10.475759

**Authors:** James W. Webber, Kevin Elias

## Abstract

**Background:** Cancer identification is generally framed as binary classification, normally discrimination of a control group from a single cancer group. However, such models lack any cancer-specific information, as they are only trained on one cancer type. The models fail to account for competing cancer risks. For example, an ostensibly healthy individual may have any number of different cancer types, and a tumor may originate from one of several primary sites. Pan-cancer evaluation requires a model trained on multiple cancer types, and controls, simultaneously, so that a physician can be directed to the correct area of the body for further testing.

**Methods:** We introduce novel neural network models to address multi-cancer classification problems across several data types commonly applied in cancer prediction, including circulating miRNA expression, protein, and mRNA. In particular, we present an analysis of neural network depth and complexity, and investigate how this relates to classification performance. Comparisons of our models with state-of-the-art neural networks from the literature are also presented.

**Results:** Our analysis evidences that shallow, feed-forward neural net architectures offer greater performance when compared to more complex deep feed-forward, Convolutional Neural Network (CNN), and Graph CNN (GCNN) architectures considered in the literature.

**Conclusion:** The results show that multiple cancers and controls can be classified accurately using the proposed models, across a range of expression technologies in cancer prediction.

**Impact:** This study addresses the important problem of pan-cancer classification, which is often overlooked in the literature. The promising results highlight the urgency for further research.

## 1. Background

Transcriptome profiling has become a powerful tool in diagnosis of cancer [24, 21, 18, 13, 14, 2, 8, 25]. Typically, in such publications, the focus is binary classification, whereby a cancer cohort of interest (e.g., pancreatic cancer [14] or breast cancer [2]) is distinguished from a control group (e.g., healthy patients [13] or a mixture of healthy patients and those with benign tumors [8]). Binary classification models are not sufficient to distinguish between different cancer types, however, as they are trained only on samples from one cancer cohort. For example, let us refer to figure 1. Here, we present synthetic data, where the samples are divided into three cohorts, namely controls, breast cancer, and ovarian cancer. In this example, a binary classifier trained to separate a single cancer type from the control group, e.g., controls vs breast cancer, is sufficient only to determine if a patient is at high risk of cancer or not, and is not suitable to distinguish specifically between breast and ovarian cancer. For visualization, samples on the upper-left side of decision boundary 1 in figure 1, would be classified (using an appropriate classifier such as a linear Support Vector Machine (SVM)) as having cancer, and vice-versa. If all samples are considered in the training, however, we can more effectively distinguish between cancer types, and controls simultaneously. In this case, a multi-classification model (e.g., SVM) has access also to decision boundary 2 of figure 1, and we can separate the sample space into three regions, one for each cohort. Thus, training a classification model on multiple cancer groups is desired, so a physician can be directed toward specific areas of the body for further testing.

**Figure 1.**
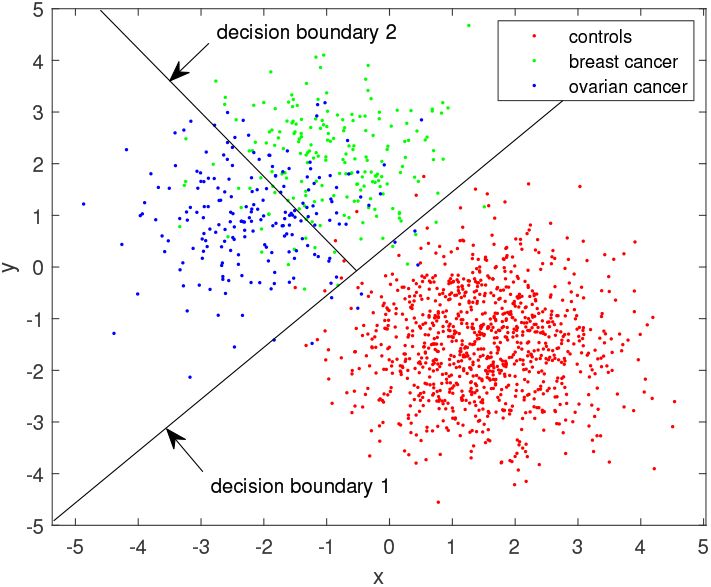
Example, synthetic multi-class data with decision boundaries.

Over the last decade, neural networks have remained a central subject of study in machine learning, with applications including image and pattern recognition [20, 3, 4, 12, 11], sensor data analysis [10], finance [9], and social network filtering [19], to name but a few. Neural networks have also been applied successfully in cancer prediction and analysis of expression data [8, 28, 26, 23]. For example, in [8], the authors employ a deep, feed-forward neural network to classify ovarian cancer patients from a control group. A comparison with a range of common classification models (e.g., SVM, Lasso, logistic regression) is presented, and the neural network is shown to offer optimal performance. In [28, 26], neural networks are applied to feature selection, and used to identify disease-related miRNA. Classification of multiple cancers using neural networks is also considered in the literature [1, 16, 17]. For example, in [16], several CNN architectures, including 1-D CNN, 2-D CNN, and hybrid CNN, are proposed to classify mRNA expression data extracted from tissue samples. The hybrid CNN takes 1-D convolutions along both image axes simultaneously (i.e., in the *x* and *y* directions), to extract features, before feeding into a dense layer for classification. The hybrid model is shown to offer optimal performance on mRNA data, when compared to conventional 1-D and 2-D CNN architectures. In [17], the authors propose a number of GCNN architectures. GCNN’s use a connectivity graph to provide further information to the network on the relation between expression variables (e.g., using a correlation matrix). Specifically, GCNN’s perform convolution among the connected variables as described by the graph Laplacian, in contrast to conventional n-D CNN’s which convolve on fixed n-D Euclidean neighborhoods (grids). The CNN and GCNN models of [16, 17] are shown to offer similar levels of performance on mRNA expression data (both papers consider the same data set).

Here, we propose novel, data-driven neural network architectures, of varying depth, which can be applied to classify multiple cancer types, and controls simultaneously across several expression technologies of interest in cancer prediction or classification (e.g., protein, miRNA, mRNA). The technique is compared against other neural network architectures proposed in the multi-cancer prediction literature. Specifically, the work of [1, 16, 17] is a main point of comparison for this study. In addition, we present an analysis of neural network complexity, with particular focus on depth, and investigate the relation to classification performance. The results show that the proposed models can classify multiple cancers and controls simultaneously, and accurately (with AUC exceeding 95%), across a range of expression technologies in cancer prediction. We discover that shallow architectures (e.g., of feed-forward type) are most optimal for multi-classification of expression data, when compared to deeper architectures. In all experiments conducted here, the proposed models are shown to offer optimal performance in terms of classification accuracy and mean sensitivity, across all data types considered, when compared to those of the literature (e.g., CNN and GCNN).

The remainder of this paper is organized as follows. In section 2 we present our results, and give a comparison between the proposed models and those of the literature. In section 3, we discuss our findings, and give our analyses as to the relation between model complexity and performance, based on the results presented. To finish, we discuss possible extensions of our work in section 4 and give our conclusions.

## 2. Results

In this section, we assess the performance of our methods on four, large expression array data sets and measurement technologies, namely miRNA [24, 21, 18, 25] (blood draw), mRNA [6] (tissue sample), and protein [15] (tissue sample). For more details on this data, see section A.4 in the appendix. We propose four, feed-forward neural network architectures, denoted NNi, for 1 ≤ *i* ≤ 4, where the index *i* denotes the depth. For example, NN2 denotes a 2-layer neural network. The proposed models are compared against other nonlinear models, namely *K*-nearest neighbors (KNN), as baseline, SVM [27], Deep (7-layer) Neural Network (DNN) [1], 1-D, 2-D and hybrid CNN models [16], and GCNN [17]. For more details on the implementation of these methods, see the methods section in appendix A.

### 2.1. miRNA expression results

Here we present our results on the miRNA expression data of [24, 21, 18, 25], collected from Japanese patients. This data and its properties are discussed in detail in point (1) of section A.4. See figure 2. In figures 2a2c, we have presented plots of the accuracy (ACC), Matthew’s Correlation Coefficient (MCC) [27], and the mean AUC, *F*_1_ score, sensitivity (Sens), specificity (Spec), and precision (Prec) over all 18 classes, for each method considered. See section A.3 for the definitions of these performance metrics. The ACC and MCC scores are included as part of figure 2a, and are labeled in the figure legend. Here CNN1, CNN2, and HCNN refer to the 1-D CNN, 2-D CNN, and Hybrid CNN models of [16], respectively. The remaining models are denoted as describes at the start of this section.

**Figure 2.**
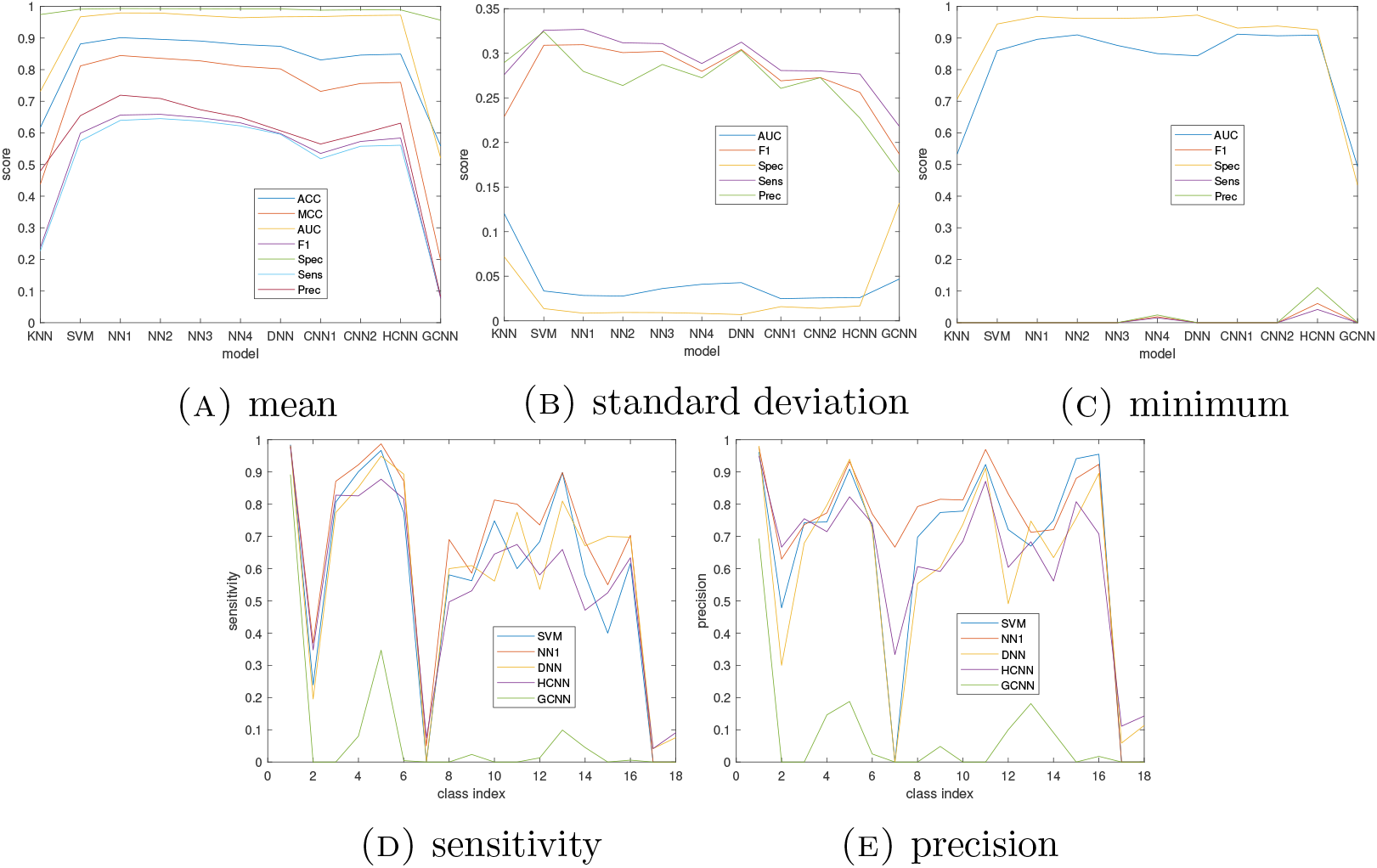
Japan data results. (A)-(C) - plots showing mean, standard deviation, and minimum scores over all 18 classes, for each method considered. The method is given on the x axis, and the metric in the figure legend. (D)-(E) - Plots showing sensitivity (recall) and precision scores for each class, for five selected methods as discussed in the main text. Here the class index is given on the x axis, and the method in the figure legend.

In figures 2d–2e, we present plots showing how the sensitivity (recall) and precision varies across classes, as is done in [16, 17], for five selected methods. We show only five out of the eleven methods considered so that the reader can more clearly compare the scores. The five chosen methods each correspond to one publication, and we have selected the best performing method (in terms of ACC) from each paper to show in figures 2d and 2e. For example, HCNN, in this case, offered the best accuracy out of CNN1, CNN2, and HCNN, proposed in [16] (see figure 2a), and was thus highlighted in figures 2d and 2e. See section A.5 in the appendix for the relations between the indices on the x axis of figures 2d and 2e, and the class (e.g., index 1, or class 1 corresponds to the control group).

Upon analysis of figure 2a, NN1 (i.e., the proposed 1-layer neural network architecture) offers the best mean score across all metrics considered, and out of all models compared against. NNi, for all 1 ≤ *i* ≤ 4, SVM, DNN, and HCNN are competitive in this regard, however. The model labeling on the *x* axis of figures 2a–2c, roughly speaking, is in order of model complexity, starting with a simple KNN approach, and then moving onto shallow, deep, convolutional, and graph convolutional neural network architectures. The mean accuracy peaks at the shallow neural net complexity level, with NN1 and NN2 offering the most competitive performance, closely followed by SVM, DNN, and HCNN. NN3 and NN4 offer no improvement in the performance, when compared to their more shallow counterparts, NN1 and NN2. The simplest approach, KNN, lacks the expressivity to describe the nonlinear separations in high-dimensional expression data, and the accuracy suffers as a result. GCNN underperforms when compared to other CNN architectures, and the graph connectivity layer employed in [17] is ineffective in extracting meaningful features from the miRNA.

The standard deviation scores, among the most competitive methods in terms of mean score (i.e., NN1-4, SVM, DNN, HCNN), are competitive, indicating a similar level of consistency. As illustrated in the sensitivity and precision plots of figures 2d and 2e, NN1 offers the best sensitivity across all classes, with the exception of class 15 (prostate cancer), where DNN is most optimal. The precision scores offered by NN1, are, for the majority of classes, greater than the methods of the literature, particularly with regards to class 7, which corresponds to biliary tract cancer. Across all methods considered, we notice a significant drop in the sensitivity scores for three classes, namely class 2 (hepatitis), class 7 (biliary tract cancer), class 17 (benign ovarian disease), and class 18 (benign ovarian tumor). To facilitate this conversation, we provide t-distributed Stochastic Neighbor Embedding (TSNE) [22] plots of the Japanese miRNA data in figure 3, so the reader can visualize how the different classes separate. In figure 3a, all samples are plotted, and we have highlighted some key populations (e.g., cancers). In figure 3b, we plot BTC samples vs other cancers. This is the left-hand cluster of figure 3a with BTC samples highlighted. The BTC samples do not appear to cluster together, as, for example, do the low-risk controls, and are scattered among the other cancers. The number of BTC samples is low (40), when compared to other cancers (e.g., there are 373 ovarian cancer samples), also. Thus, NN1 (and all other models considered) fails to accurately classify the BTC samples and the class sensitivity is close to zero. This could be a sample size issue, given the relatively small sample size for BTC. That is, the number of BTC samples is likely insufficient to observe clustering. We may start to see clustering among the BTC cohort, as more samples become available. Further data collection is needed, however, to confirm this hypothesis.

**Figure 3.**
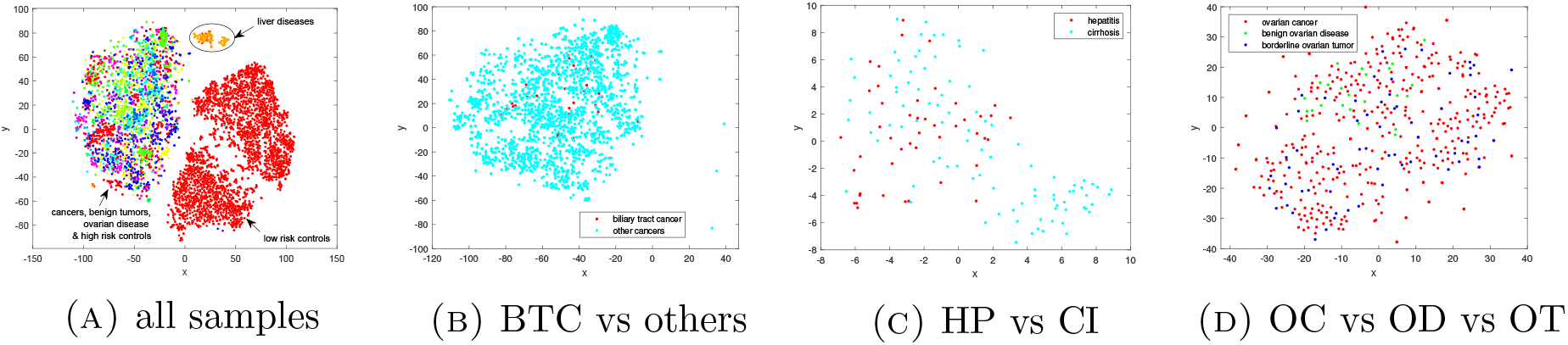
TSNE plots of Japanese miRNA data. (A) - all samples with some key groupings (e.g., low risk controls) highlighted. (B) - Biliary Tract Cancer (BTC) patients vs other cancer patients. (C) - hepatitis (HP) vs cirrhosis (CI). (D) - Ovarian Cancer (OC) vs benign Ovarian Disease (OD) vs benign Ovarian Tumor (OT).

In figure 3c, we plot the liver disease samples, which are highlighted in figure 3a in orange. The plots show significant mixing among HP and CI samples. Given the mixing effect, and since CI is the larger class, with 93 CI samples vs 46 HP samples, the neural network classifies the majority of HP patients as CI, thus yielding low HP sensitivity. We see a similar mixing effect in the OC vs OD vs OT plots of figure 3d. In this case, the neural network classifies all patients with benign ovarian disease and benign ovarian tumors as having ovarian cancer, again given also the discrepancies in sample size, with 373 OC, 24 OD, and 66 OT patients. This goes to show that, while we can accurately distinguish between more general disease groupings (e.g., an ovarian abnormality vs a pancreatic abnormality), the specific issue (e.g., ovarian cancer, ovarian disease etc.) is difficult to identify with a neural network model. This is where a physician would come in. That is, once we have identified the area of concern (e.g., the ovaries) using a neural network and miRNA expression, a physician can conduct further tests (e.g., imaging, biopsy) to ascertain the specific issue.

### 2.2. protein data results

Here we present our results on the protein data collected as part of The Cancer Proteome Atlas (TCPA) [15]. As in section 2.1, in figure 4 we display plots of the mean, standard deviation, and minimum classification scores, and show how the sensitivity and precision varies across classes.

**Figure 4.**
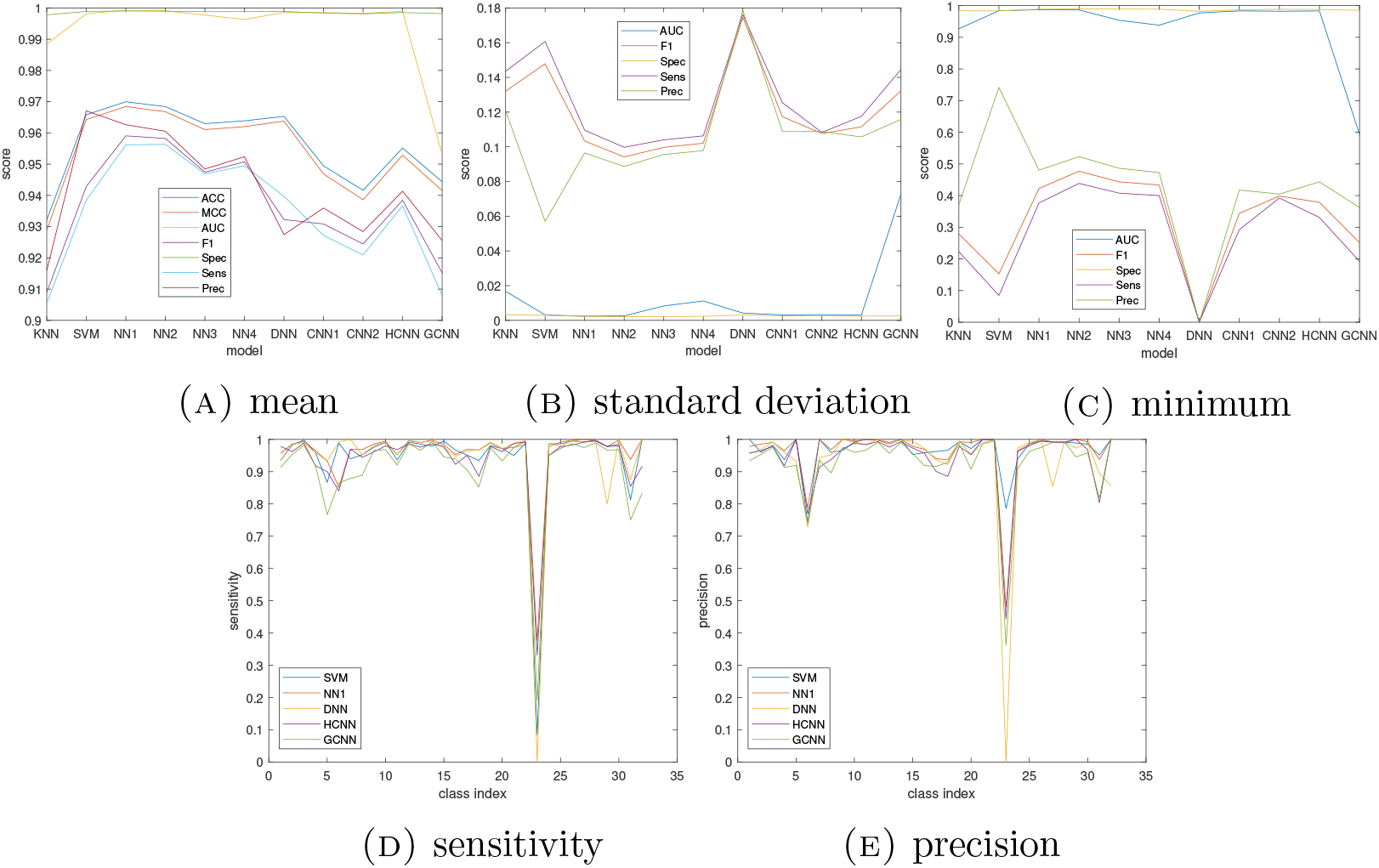
TCPA protein data results. (A)-(C) - plots showing mean, standard deviation, and minimum scores over 32 classes, for each method considered. The method is given on the x axis, and the metric in the figure legend. (D)-(E) - Plots showing sensitivity (recall) and precision scores for each class, for five selected methods as discussed in the main text. Here the class index is given on the x axis, and the method in the figure legend.

In this example, NN1 offers the best mean score across all metrics, with the exception of precision, where SVM is most optimal. See figure 4a. As was discovered also in section 2.1 on miRNA data, KNN and GCNN perform relatively poorly when compared to the other methods considered, although the accuracy offered by KNN and GCNN is good and exceeding 90%. We notice a peaking effect in the mean score curves of figure 4a at the low-level neural network complexity, with NN1 and NN2 offering the most competitive performance. This is also consistent with the findings of section 2.1. We see a reduction in ACC, MCC, mean sensitivity and precision with increasing neural network complexity, and when the depth of the neural network exceeds two layers. Out of CNN1, CNN2, and HCNN, i.e., the models proposed in [16], HCNN offers the best mean performance across all metrics considered. The same was true in section 2.1 on miRNA data. This is consistent with the findings of [16], where HCNN was shown to offer the best performance on mRNA data.

In figures 4b and 4c, we present plots of the standard deviation and minimum score, for each method considered. NN2 offers the most consistent performance, with the smallest standard deviation and greatest minimum, for all metrics considered, with the exception of precision, for which SVM offers the best consistency.The performance offered by SVM, and NN1-4 is competitive, in terms of standard deviation and minimum score across classes. CNN1, CNN2, HCNN, and GCNN are also reasonably competitive in this regard. In figures 4d and 4e, while, for the majority of classes, the sensitivity and precision is consistently high, and generally exceeding 90%, particularly for the more competitive methods, such as NN1 and SVM, we notice a significant drop in the sensitivity and precision at class 25, which corresponds to rectum adenocarcinoma (READ). NN1 maintains the best sensitivity for READ, and SVM, the best precision. The standard deviation and minimum curves of figures 4b and 4c show large jumps in sensitivity at KNN, SVM, and DNN. These models offer close to zero sensitivity for the READ class, which acts as an outlier in the standard deviation calculations, and creates the observed spiking effect.

To help explain the sharp decline in READ accuracy, refer to figure 9 in appendix A.9, where we have presented TSNE plots of the TCPA data. In figure 9c, we plot all samples. We see a strong clustering effect among the majority of the 32 cancers, with the exception of READ and colon adenocarcinoma (COAD) samples, which appear mixed. See figure 9d. The number of COAD samples far outweighs the READ samples, with 357 COAD and 130 READ samples, giving greater prior to a COAD classification. Thus, NN1 (and the other models considered here) classifies the majority of READ samples as COAD. So, while NN1 cannot accurately distinguish between cancerous tissue drawn from the colon, more generally, and the rectum (i.e., the last 12cm of the colon), it can distinguish with high sensitivity cancerous tissues from different parts of the body. Overall, the ability of NN1 (and all other models considered) to distinguish different cancers is much improved when using protein data drawn from tissue, when compared to miRNA expression sequenced from blood, as indicated by the classification performance and TSNE plots.

### 2.3. mRNA data results

In this section, we present our results on the mRNA expression data collected as part of The Cancer Genome Atlas (TCGA) [6]. See figure 5 for plots of our results, presented in the same way as in sections 2.1 and 2.2.

**Figure 5.**
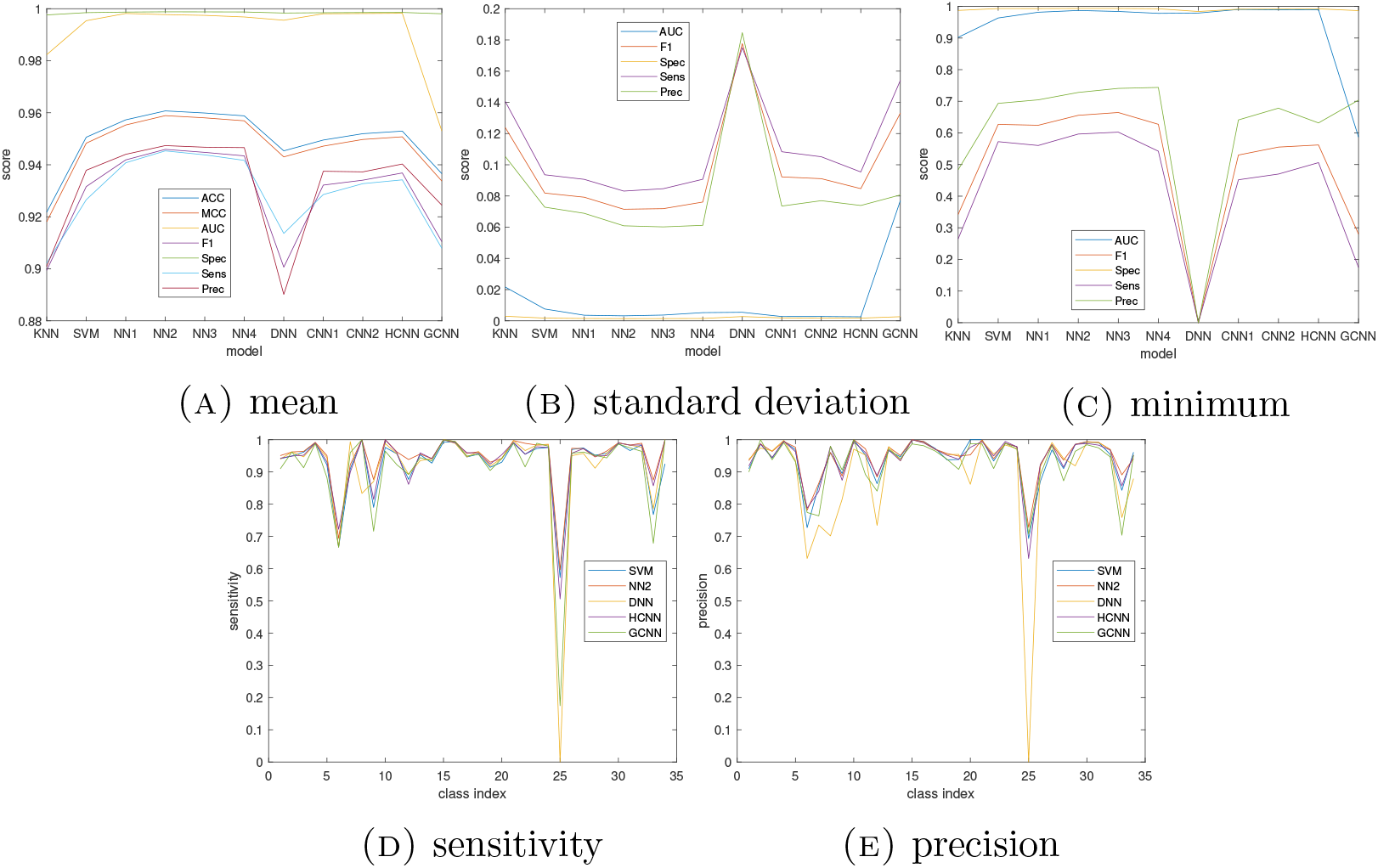
TCGA mRNA data results. (A)-(C) - plots showing mean, standard deviation, and minimum scores over 34 classes, for each method considered. The method is given on the *x* axis, and the metric in the figure legend. (D)-(E) - Plots showing sensitivity (recall) and precision scores for each class, for five selected methods as discussed in the main text. Here the class index is given on the *x* axis, and the method in the figure legend.

In this example, NN2 offers the best mean and standard deviation performance across all metrics considered, with minimum scores that are competitive with SVM, NN1, NN3, and NN4. See figures 5a–5c. NN2 also offers greater than or equal sensitivity and precision, when compared to the methods of the literature, for the majority classes. See figures 5d and 5e. In [16] and [17], the authors consider the same data set as in this section (i.e., TCGA data). In light of this, we wish to explain the slight discrepancies in the accuracy scores reported here and those of [16, 17]. In [16], 5-fold Cross Validation (CV) is used to test model accuracy, and in [17], a hold out set is used. In this paper, we use 10-fold CV, and the train-test splits are kept fixed throughout, for fairness. Hence, the train-test splits used here are different to that of [16, 17], thus leading to a slight change in accuracy. For example, the classification accuracy (ACC) reported here for HCNN with 10-fold CV is 95.3%, which is slightly higher than the 95.0 ± 0.1% score reported in [16].

As was discovered in section 2.2, with protein data, we notice a significant drop in the class sensitivity and precision at class 25, which corresponds to READ. We see drop in sensitivity at class 6, also, which corresponds to cholangiocarcinoma (CHOL), although the reduction in sensitivity is less significant. The drop in READ sensitivity is due to mixing with COAD samples, and discrepancies in class size between READ and COAD. Such effects were also observed in our analysis of protein data in section 2.2. The drop in CHOL sensitivity is due to mixing with liver hepatocellular carcinoma (LIHC) samples, although the overlap is less pronounced. The READ vs COAD, and CHOL vs LIHC mixing effects are discussed and highlighted in [16], and TSNE plots are provided, and thus we do not discuss the overlap in great length. For completeness, we provide TSNE plots of the TCGA data here also in the appendix. See appendix A.9. NN2 offers the best sensitivity and precision for the READ class, and is competitive with SVM and HCNN. DNN, in this case, grouped all READ samples into the COAD class, yielding zero sensitivity, which explains also the reduced performance of DNN, in terms of mean, standard deviation, and minimum accuracy, when compared to the other methods of the literature and the proposed.

## 3. Discussion

In this paper, we introduced novel neural network models for the early detection of multiple cancers, which are applicable to a variety of expression technologies, including protein, miRNA, and mRNA. The models presented were shown to outperform those of the literature [27, 16, 1, 17]. Upon analysis of the model complexity and neural network depth, we discovered that the more shallow neural network architectures (e.g., NN1 and NN2) fared better when compared to deeper, more complex architectures such as DNN [1], CNN [16] or GCNN [17]. In particular, the plots of figures 2a, 4a, and 5a, which show how the mean classification metrics (such as ACC, MCC, F_1_ score,and AUC) vary with model complexity, peaked at the shallow neural network level, and NN1 and NN2 (i.e., the 1 and 2-layer neural network architectures proposed here) offered the best performance in terms of ACC, MCC and F_1_ score, for all data types considered. All methods compared against were competitive in terms of mean AUC. With increasing depth, we either saw no increase in mean accuracy (e.g., as in figure 5a) or a slight decrease (e.g., as in figure 4a), and the deep feed-forward models NN3, NN4 and DNN (7-layers) [1] offered no improvement in accuracy when compared to NN1 and NN2. The degradation in accuracy was often more pronounced for convolutional models such as CNN1, CNN2, and HCNN, and more so for GCNN. To help explain this, let us refer to figure 6. In sub-figures 6b and 6d, we show classical examples of data where CNN type models offer state-of-the-art classification performance. For example, figure 6b in an image taken from the MNIST database [7]. 2-D CNN’s are proven to offer optimal classification performance on MNIST data [5]. The data in figures 6b and 6d has a clearly visible structure (e.g., the shape, edges, and size of the handwritten four), and a convolution filter can extract valuable features (e.g., an edge map of the handwritten four, a wavelet decomposition of the airline passenger time series) from such information. It is the aim of the CNN training to learn such a convolution filter. The miRNA expression data does not share such structure. See figures 6a and 6c. Here we have shown a sample taken from the Japanese data set [24, 21, 18, 25], and reshaped the sample to form an image in figure 6a. See section A.6 in the appendix for example images and plots of mRNA and protein data. A lack of structure is apparent also for this data. Thus, expression data appears inconsistent with the types of applications where CNN type models are known to perform well, which may help to explain the reduction in CNN accuracy observed in the experiments conducted here.

**Figure 6.**
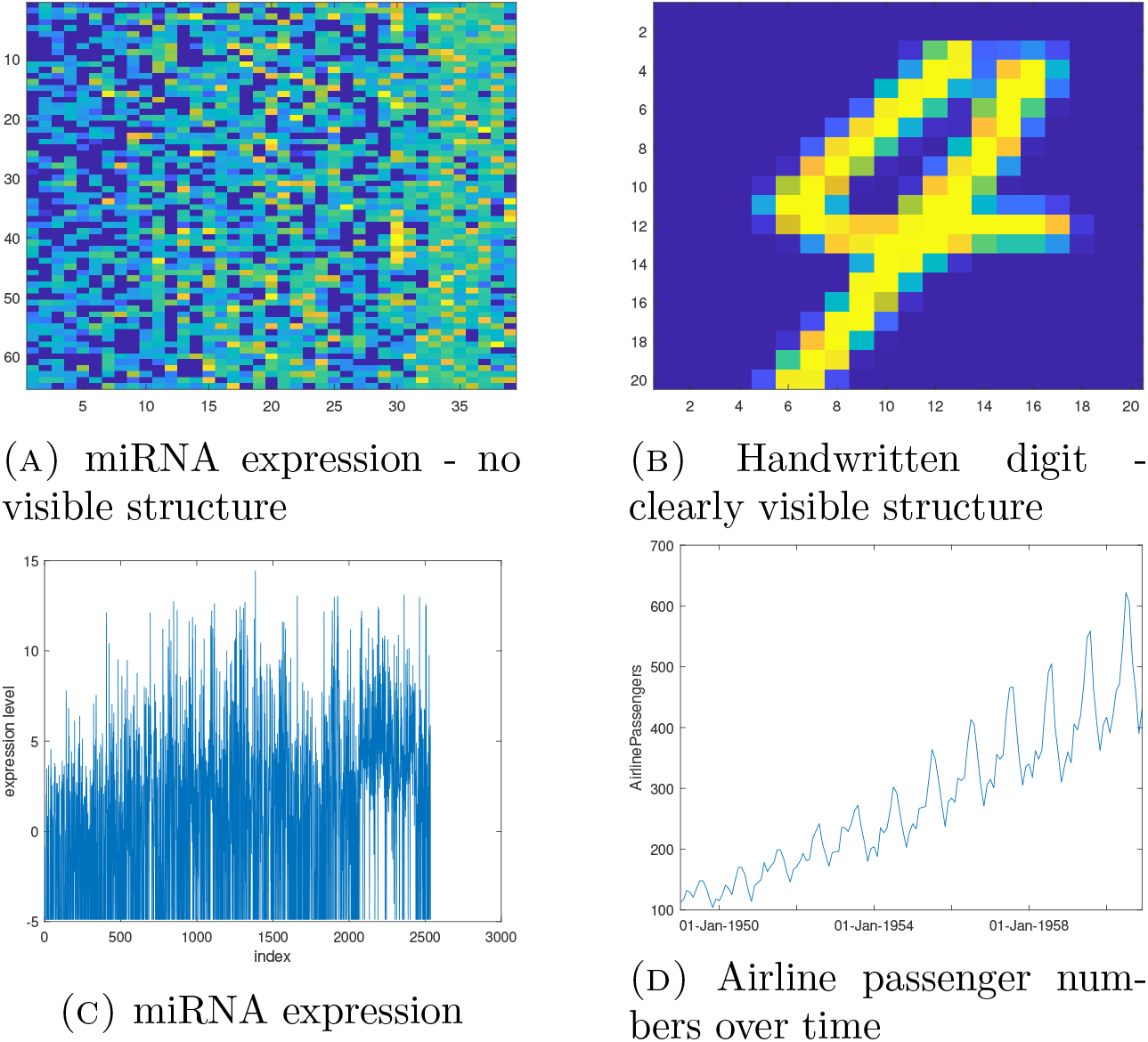
Top row - example images of miRNA expression and handwritten image data. Bottom row - example plots of miRNA expression, and airline passenger numbers over time.

Let us refer back to the synthetic example discussed in the introduction in figure 1. In the TSNE plots presented in figure 3a, we showed real miRNA data examples of the clustering effect among cancers and controls, although the separations were not so clear and linear as in figure 1. In practice, the separation among cancers and controls in high-dimensional expression space is nonlinear. Neural networks are well-known to be effective in untangling such nonlinearities in high-dimensional data. However, it is not always clear how one should decide upon a specific architecture for a given problem. This analysis provides insight to this end in regard to cancer prediction applications. Based on the results presented here, we find that shallow, feed-forward architectures offer optimal performance, for the problem of classifying multiple cancers using expression data.

## 4. Conclusions and further work

This study provides a simple, robust approach to multi-cancer classification, applicable to several commonly applied expression technologies in cancer prediction (e.g., miRNA, mRNA, and protein), which is shown to outperform alternative neural network techniques previously introduced in the literature. Upon investigation of the neural network complexity, with particular regard to depth, shallow models were shown to offer optimal accuracy, when compared more complex models, such as DNN [1] and CNN [16].

In future work, we aim to investigate further the utility of neural networks in multicancer prediction, and expand on the current study, which focused on feed-forward, CNN, DNN, and GCNN type architectures from the literature. In particular, we aim to pursue ideas in manifold learning to better understand and visualize the nonlinear boundaries between different cancers and controls in high-dimensional expression space, and to inform our decisions on network selection.

## 5. Declarations

Here we give our declarations.

### 5.1. Ethics approval and consent to participate

No new human or animal data is presented here.

### 5.2. Consent for publication

There are no issues regarding consent for publication.

### 5.3. Availability of data and materials

All real data sets considered here are publicly available online. See section A.4 for more details. The neural network code and the synthetic data discussed in section 1 is available from the authors upon reasonable request.

### 5.6. Authors’ contributions

JWW developed the neural network models and the original idea. The experiments were conducted by JWW. Analyses of results by JWW and KE. KE was as major contributor in writing the manuscript, and provided expert insight from a medical background needed to communicate this work to a medical audience.

## Appendix A. Methods

Here we describe in detail our approach, the methods from the literature compared against, the validation methods used for testing, and selection of hyperparameters.

### A.1. The proposed architecture

In this section, we explain the details of the proposed neural network architectures, NN1-4. NN1-4 are feed-forward type and fully connected. The activation functions considered were Rectified Linear Unit (ReLU), Exponential Linear Unit (ELU), sigmoid, hyperbolic tangent, and softplus, and the layer sizes (i.e., number of neurons) were chosen from {64,128, 256}. To select the activation function and number of neurons at each layer, we took a data-driven approach. Specifically, we employed nested cross validation as described in section A.8. The networks were trained in Tensorflow v-2.6 using RMSprop. The code to implement all network training and validation is available upon reasonable request to the authors.

### A.2. Methods for comparison

The proposed architectures are compared against *K*-Nearest Neighbors (KNN), Support Vector Machine (SVM), and the neural network models proposed in [1, 16, 17]. An SVM model is employed in [27] to classify ten cancer types simultaneously. Our SVM model is similar to that of [27], only trained in a different way. As no exact code is provided in [27], we fit our SVM using the inbuilt Matlab (v-2020b) function “fitcecoc”. The KNN model is trained using the inbuilt Matlab code, “fitcknn”, with a Euclidean distance metric. The number of neighbors (K) was chosen as described in section A.8.

The deep neural network model (DNN) of section 2 was constructed as described in [1], and trained using algorithms as specified in https://github.com/sipark5340/PathDeep. This link was provided to us by the authors of [1]. The CNN models, CNN1, CNN2, and HCNN, presented were trained using the code provided in [16]. In the Github link provided by the authors (https://github.com/chenlabgccri/CancerTypePrediction), there are two models provided for each network type. For example, two 1-D CNN’s are proposed, one which is designed for classification of cancerous and normal tissue, and one exclusively for cancerous tissue. In all examples conducted here, and for each model of [16], we presented the network which offered the best classification accuracy (ACC).

The graph convolutional neural networks considered here (model GCNN from section 2) were trained using the code of [17]. For the TCGA data example (in section 2.3), we use the exact code provided in the Github link https://github.com/RicardoRamirez2020/GCN_Cancer, provided in [17]. In [17], the authors consider the same TCGA data [6] as in this paper, to test their GCNN model. The authors provide a connectivity graph based on Protein-Protein Interaction (PPI). A connectivity graph based on correlation matrix is also provided. The PPI model is shown to offer optimal performance on TCGA data in [17], and hence why we compare against PPI here in the TCGA example of section 2.3. In sections 2.1 and 2.2, since no PPI graph is provided in [17] for miRNA or TCPA data, we construct a connectivity graph using a correlation matrix, as described in [17]. In the results of [17], the correlation matrix and PPI models are shown to be highly competitive, so we do not expect much loss in accuracy for lack of a PPI graph in these examples. We also include singleton graph nodes, in all examples, so that no data is discarded before network training.

### A.3. Classification metrics

Here we introduce the metrics which will be used to assess the quality of the results. Let TP, FP, TN, and FN denote the number of true positives, false positives, true negatives, and false negatives, respectively. Then, we consider the following classification metrics in our comparisons:

1. The classification accuracy (denoted by “ACC”)

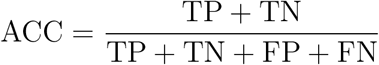
2. The *F***i** score

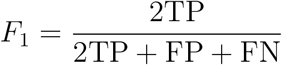
3. The True Positive Rate (TPR, also known as sensitivity or recall)

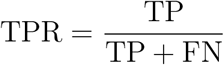
4. The True Negative Rate (TNR, also known as specificity)

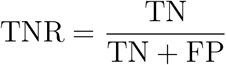
5. Positive Predictive Value (PPV), also known as precision

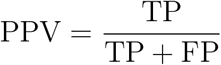
6. Matthew’s Correlation Coefficient (MCC). This is defined on page 6 of [27] for multi-class problems.

We also report the Area Under the receiver operator characteristic Curve (AUC) values. All metrics reported here take values between 0 and 1. A value closer to 1 indicates a better performance, and vice versa.

### A.4. Data

We consider the following four data sets from the literature to assess model performance:

1. Serum miRNA expression data of [24, 21, 18, 25] collected from 2460 Japanese patients. Here the authors provide expression values for 3781 control patients, and for patients with 17 different types of diseases (the case number varies with the disease), including bladder cancer, hepatocellular carcinoma, breast cancer, ovarian cancer, and hepatitis. To compile the data used here, we combine the data sets of [24, 21, 25], making sure to delete any replicate patients. In [18] expression values are provided for 79 patients with advanced breast cancer. We combine the advanced breast cancer samples of [18] with the breast cancer samples of [24, 21, 25] to form a breast cancer cohort, yielding 234 breast cancer samples in total. In [18], the number of expression values recorded is 2535, which is slightly smaller than the 2565 of [24, 21, 25]. In this study, we use the 2535 miRNA’s of [18] so that the data sets can be combined.
2. Protein data of [15], collected as part of TCPA. The data is comprised of 7694 tissue samples from 32 different cancers, which are summarized in column 3 of table 1. The data was downloaded specifically from https://www.tcpaportal.org/tcpa/download.html under the “Pan-Can 32” directory.
3. mRNA expression data of [6], collected as part of TCGA. The data is comprised of 10339 samples from 33 different cancer tissues, and 731 samples from noncancerous (normal) tissue. The 33 cancer types of TCGA include all 32 TCPA cancers, with the addition of acute myeloid leukemia (LAML). In total, there are 10339 + 731 = 11070 sampes. The data was downloaded from https://github.com/RicardoRamirez2020/GCN_Cancer, which also provides code for training GCNN models as described in [17].

**Table 1.** Class indices labeling.

| class index | miRNA [24, 21, 18, 25] | protein [15] | mRNA [6] |
| --- | --- | --- | --- |
| 1 | control | ACC | NT |
| 2 | hepatitis | BLCA | ACC |
| 3 | cirrhosis | BRCA | BLCA |
| 4 | HCC | CESC | BRCA |
| 5 | bladder cancer | CHOL | CESC |
| 6 | breast cancer | COAD | CHOL |
| 7 | BTC | DLBC | COAD |
| 8 | colorectal cancer | ESCA | DLBC |
| 9 | esophageal cancer | GBM | ESCA |
| 10 | gastric cancer | HNSC | GBM |
| 11 | glioma | KICH | HNSC |
| 12 | lung cancer | KIRC | KICH |
| 13 | ovarian cancer | KIRP | KIRC |
| 14 | pancreatic cancer | LGG | KIRP |
| 15 | prostate cancer | LIHC | LAML |
| 16 | sarcoma | LUAD | LGG |
| 17 | OD | LUSC | LIHC |
| 18 | OT | MESO | LUAD |
| 19 |  | OV | LUSC |
| 20 |  | PAAD | MESO |
| 21 |  | PCPG | OV |
| 22 |  | PRAD | PAAD |
| 23 |  | READ | PCPG |
| 24 |  | SARC | PRAD |
| 25 |  | SKCM | READ |
| 26 |  | STAD | SARC |
| 27 |  | TGCT | SKCM |
| 28 |  | THCA | STAD |
| 29 |  | THYM | TGCT |
| 30 |  | UCEC | THCA |
| 31 |  | UCS | THYM |
| 32 |  | UVM | UCEC |
| 33 |  |  | UCS |
| 34 |  |  | UVM |

### A.5. Class indices table

Table 1 describes the cohorts with which the indices on the x axis of figures 2d–2e, 4d–4e, and 5d–5e correspond. In table 1, ACC stands for AdrenoCortical Cancer, not to be confused with classification accuracy (also denoted ACC), defined in section A.3. NT stands for Normal Tissue. The remaining nomenclature for the TCGA/TCPA data is defined in [17, page 14]. In the second column of table 1, HCC stands for HepatoCellular Carcinoma, BTC stands for Biliary Tract Cancer, OD for benign Ovarian diseases, and OT for benign Ovarian Tumor. The data sets and expression technologies (with references) are given on the first row and the class indices in the first column.

### A.6. Example data plots and images

Here we present example images and plots of mRNA [6] and protein [15] data. An example for miRNA data is provided in the main text in section 3. In figures 7 and 8, we show example mRNA and protein data, as in section 3. The images and plots indicate lack of structure, as is, for example, seen in handwritten image or time series data (see figures 6b and 6d), where CNN type models are known to perform well. This might help explain the drop in mean accuracy observed in figures 2a, 4a, and 5a, when going from feed-forward to CNN type architectures.

**Figure 7.**
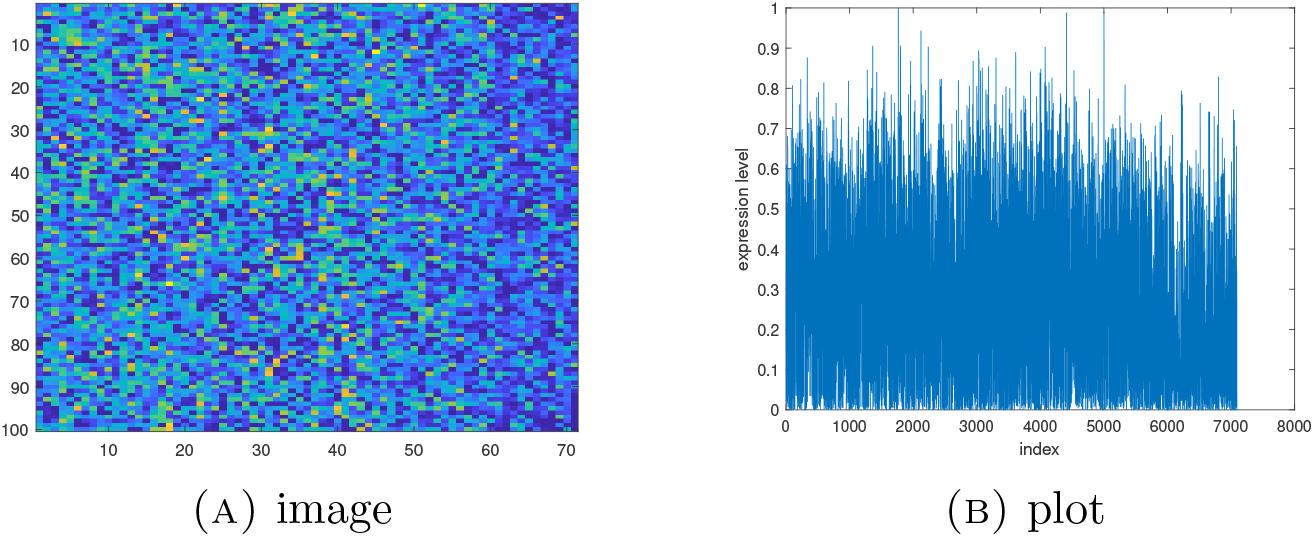
Example image and plot of mRNA TCGA data.

**Figure 8.**
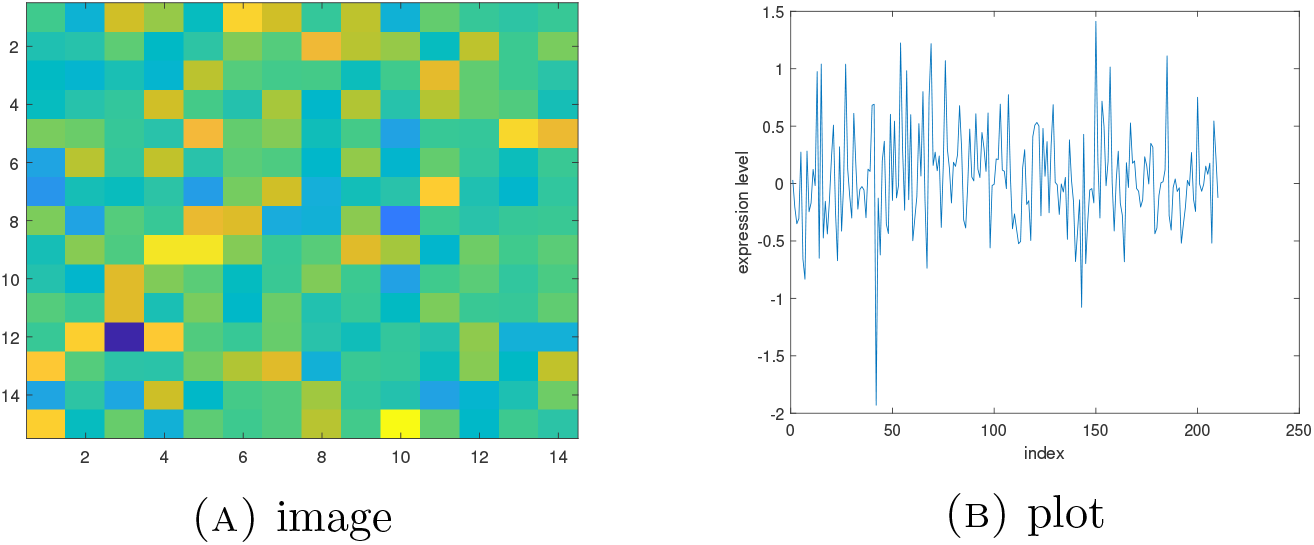
Example image and plot of TCPA protein data.

### A.7. Validation methods

All methods compared against were validated using 10-fold cross validation, for all data sets considered. The specific train-test splits were kept fixed throughout, for fairness.

### A.8. Hyperparameter selection and normalization

For NN1-4, SVM, KNN, and DNN, all data was normalized to the cube [-1,1]^p^, where *p* is the number of variables, and centered with mean zero. For the remaining models, i.e., CNN1, CNN2, HCNN, and GCNN, the data was normalized as described in the code provided by the authors in [16, 17]. That is, we applied the exact code of [16, 17] so as to accurately represent the authors work. The hyperparameters for NN1-4 (e.g., number of neurons, layer activations) were selected by nested cross validation. Specifically, the model selection was performed by multiple hold-out cross validation, using only the training data, to be sure of no optimism due to model selection bias. The number of neighbors for KNN was selected in the same way. DNN, CNN1-2, HCNN, and GCNN are fixed architectures from the literature, and thus require no further hyperparameter tuning.

### A.9. Additional TSNE plots

Here we present TSNE plots of the TCGA mRNA data and TCPA protein data, which were considered in sections 2.3 and 2.2. Such plots are provided in [16, page 9], but we include them here also for completeness, and for self-containment of the article. See figure 9 for TSNE plots of TCGA and TCPA data. In sub-figures 9a and 9c, we plot all samples, and the different cancer/noncancer tissues are highlighted in different colors. We see good separation among the different classes for the most part, although the READ and COAD samples appear mixed. See sub-figures 9b and 9d, where we have zoomed in on the COAD and READ samples to highlight the mixing effect.

**Figure 9.**
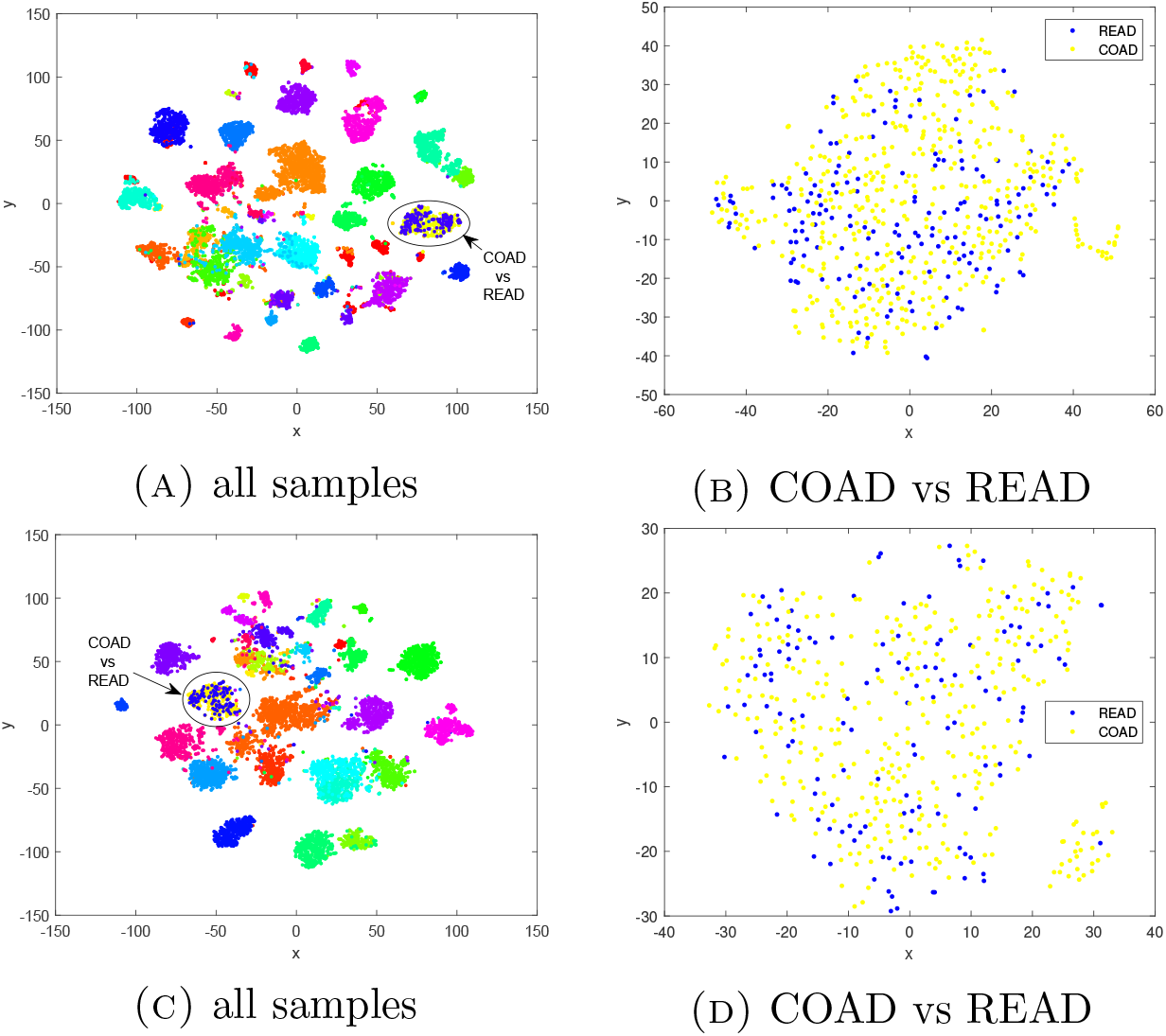
TSNE plots of TCGA [6] (top row) and TPCA 2.2 (bottom row) data. Left – all samples with READ vs COAD highlighted. Right – shows mixing of COAD and READ samples by zooming in on the circled regions of the left-hand plots.

